# Capturing continuous spectrotemporal pupil–brain coupling with simultaneous wearable EEG/Eye tracking

**DOI:** 10.64898/2026.11.08.765166

**Authors:** Tongxin Liu, Sonja A. Kotz, Michael Schwartze, Antonio Criscuolo

**Affiliations:** Department of Neuropsychology & Psychopharmacology, Faculty of Psychology and Neuroscience, Maastricht University, 6200 MD, Maastricht, the Netherlands; Department of Neurology, Max Planck Institute for Human Cognitive and Brain Sciences, 04103 Leipzig, Germany; Institute of Medical Psychology, University of Lübeck, 23562 Lübeck, Germany; Max Planck Institute for Biological Cybernetics, 72076 Tübingen, Germany

**Keywords:** Pupillometry, EEG, Resting state, Pupil–brain coupling, Lightweight wearable devices, Granger causality

## Abstract

Spontaneous fluctuations in pupil diameter are closely linked to neural dynamics, yet most evidence for pupil–brain coupling has been obtained in conventional laboratory settings. We therefore asked whether the spectrotemporal and directional organization of such coupling can be captured with wearable systems to test their use in naturalistic settings. Using combined lightweight mobile EEG (MUSE) and eye tracking (Tobii Pro Glasses 3) devices, we investigated resting-state pupil–brain coupling and explored interoceptive attention to determine whether the core organizational principles observed with stationary equipment are detectable within and preserved across continuous recording states. Cross-correlation, mutual information, and Granger causality analyses revealed that pupil–brain coupling displayed clear frequency-dependent organization. Lower EEG frequencies (2-12Hz) preceded pupil dynamics by roughly 764ms, while higher frequencies showed a distinct coupling range. Granger-causality analysis further indicated predominantly EEG-to-pupil directional coupling. Similar observations were made during interoception. Overall, these findings align well and extend previous evidence on spontaneous pupil–brain coupling, demonstrating that lightweight and movable wearable devices are suitable to characterize spectrotemporal and directional pupil– brain interactions. In turn, these results encourage future fundamental and translational research in naturalistic environments.

## 1. Introduction

Pupil diameter is increasingly used as a directly accessible physiological index of continuous fluctuations in arousal and brain state (Joshi & Gold, 2020; Liu et al., 2026; Lloyd et al., 2023). Beyond responses to luminance changes and discrete cognitive events, pupil diameter also exhibits spontaneous fluctuations in the absence of overt external stimulation (Bouma & Baghuis, 1971; Yoss et al., 1970). These spontaneous pupil fluctuations (SPFs) are linked to activity within several neuromodulatory systems, including noradrenergic and cholinergic pathways (Joshi & Gold, 2020; Joshi et al., 2016; Lloyd et al., 2023; Reimer et al., 2016), as well as activity across widespread cortical regions (Lloyd et al., 2023). Consequently, SPFs are part of a broader pupil–brain framework, in which changes in pupil size provide a peripheral index of interactions between neuromodulatory activity and cortical state (Liu et al., 2026). Their non-invasive nature and ease of acquisition make SPFs particularly suitable for continuous monitoring across relatively unconstrained recording settings. An important question is whether these pupil-linked neural dynamics can also be detected using lightweight mobile recording systems. Establishing this feasibility would open opportunities to investigate pupil–brain coupling in more naturalistic and ecologically valid settings.

Prior data from animal electrophysiology and human EEG and MEG suggest that the link between spontaneous pupil fluctuations and cortical activity is strongly frequency dependent. In rodents, increased pupil size is accompanied by enhanced gamma activity and reduced low-frequency cortical activity, particularly around 2-10 Hz (McGinley et al., 2015; Reimer et al., 2014). Similarly, human studies showed that pupil-linked changes are expressed across distinct spectral ranges, including delta, alpha, beta, and gamma activity (Ceh et al., 2020; Montefusco-Siegmund et al., 2022; Pfeffer et al., 2022; Podvalny et al., 2021). Such coupling is also temporally structured. Cortical activity has been reported to precede subsequent pupil fluctuations by several hundred milliseconds, with the estimated lag ranging from approximately 200 ms in the alpha range to more than 800 ms at lower frequencies (Montefusco-Siegmund et al., 2022; Pfeffer et al., 2022), and broader MEG activity has been shown to precede rapid pupil fluctuations by around 400 ms (Podvalny et al., 2021). Together with resting-state fMRI evidence of temporally structured associations between pupil fluctuations and large-scale cortical networks (Yellin et al., 2015), these findings indicate that pupil–brain coupling is more differentiated than any broadband relationship and instead shows a systematic spectrotemporal organization.

Despite the growing evidence for a systematic spectrotemporal organization, the directionality of spontaneous pupil–brain coupling remains less understood. Temporal order alone does not establish whether preceding neural activity provides predictive information about subsequent pupil dynamics, and vice versa. Resting-state fMRI studies reported bidirectional temporal dependencies between pupil fluctuations and activity in different cortical regions (Yellin et al., 2015), but frequency-resolved evidence from electrophysiological signals remains limited. As highlighted in a recent review of the resting-state pupil–brain system, a more comprehensive characterization of the organizational principles of the pupil–brain link depends on a better understanding of when and across which frequencies pupil and neural signals covary, but also of how their coupling scales (Liu et al., 2026). A better understanding of these directional relationships could therefore complement existing evidence on the temporal and frequency-dependent organization of spontaneous pupil– brain coupling. Most previous evidence on pupil–brain coupling has been obtained using stationary EEG, MEG, or fMRI systems in conventional laboratory settings. As pupil dynamics are sensitive to environmental factors such as ambient luminance, it is possible that pupil–brain coupling may be expressed differently in naturalistic environments. Secondly, portable and low-cost EEG technologies have made (electro-)physiological recordings increasingly feasible with reduced setup requirements, budget, and higher mobility (De Vos & Debener, 2014; Debener et al., 2012; Krigolson et al., 2017). Whether the temporal and frequency-dependent organization of spontaneous pupil–brain coupling can be recovered using a lightweight, low-channel EEG system, together with eye tracking remains to be tested. Establishing this use would help determine whether pupil–brain coupling reflects a robust feature of continuous brain-body dynamics rather than a relationship that depends on highly controlled laboratory conditions. Furthermore, such an approach could facilitate the study of pupil–brain dynamics ‘in the wild’, opening opportunities for ecologically valid research with basic and translational applications.

As a step towards investigating pupil–brain coupling in more naturalistic settings, we examined whether the spectrotemporal organization of spontaneous pupil–brain coupling could be recovered using lightweight wearable EEG together with mobile eye tracking. We further extended this characterization by examining directional coupling between EEG and pupil dynamics using spectral Granger causality. Resting state was the primary condition of interest, allowing spontaneous pupil and neural fluctuations to be examined in the absence of external task demands. As a secondary condition, interoceptive attention introduced an explicit attentional constraint by directing participants toward bodily sensations. This condition allowed exploring whether the core organization observed at rest would be maintained during an internally oriented attentional state.

## 2. Methods

### 2.1 Participants

Forty-one healthy young adults (33 female; mean age = 23.56 years, SD = 5.21; range, 18–37 years) were recruited at Maastricht University. Participants reported no history of ophthalmological or neurological disorders and provided written informed consent before participation. The study was approved by the ethics committee of the University of Maastricht (OZL_255_97_07_2022).

### 2.2 Experimental procedure

The experiment was conducted in a quiet laboratory under constant ambient lighting. Participants were seated comfortably in a chair and instructed to fixate on a blue cross presented at the center of a computer screen against a gray background. They first completed the 7 min resting-state condition, during which they were instructed to remain still and maintain fixation. This was followed by the 7 min interoceptive attention condition, which required the participants to direct their attention toward their breathing and heartbeat while maintaining fixation.

### 2.3 Data acquisition

All signals were acquired using wearable and portable recording devices (Fig. 1; see Criscuolo et al. (2026), for details on experimental setup). Pupil diameter was recorded binocularly using Tobii Pro Glasses 3 (Tobii, Stockholm, Sweden; https://www.tobii.com) at a sampling rate of 50 Hz. The eye tracker was calibrated before each condition to ensure accurate pupil recording. EEG activity was recorded using a MUSE S headband (InteraXon Inc., Toronto, Canada; https://choosemuse.com) from four channels (TP9, AF7, AF8, and TP10) at a sampling rate of 250 Hz. Electrocardiographic (ECG) activity was recorded using a Polar H10 heart rate sensor (Polar Electro Oy, Kempele, Finland; https://www.polar.com). The ECG data were not included in the pupil–brain coupling analyses and were used solely during EEG preprocessing to identify and correct cardiac artifacts.

**Fig. 1.**
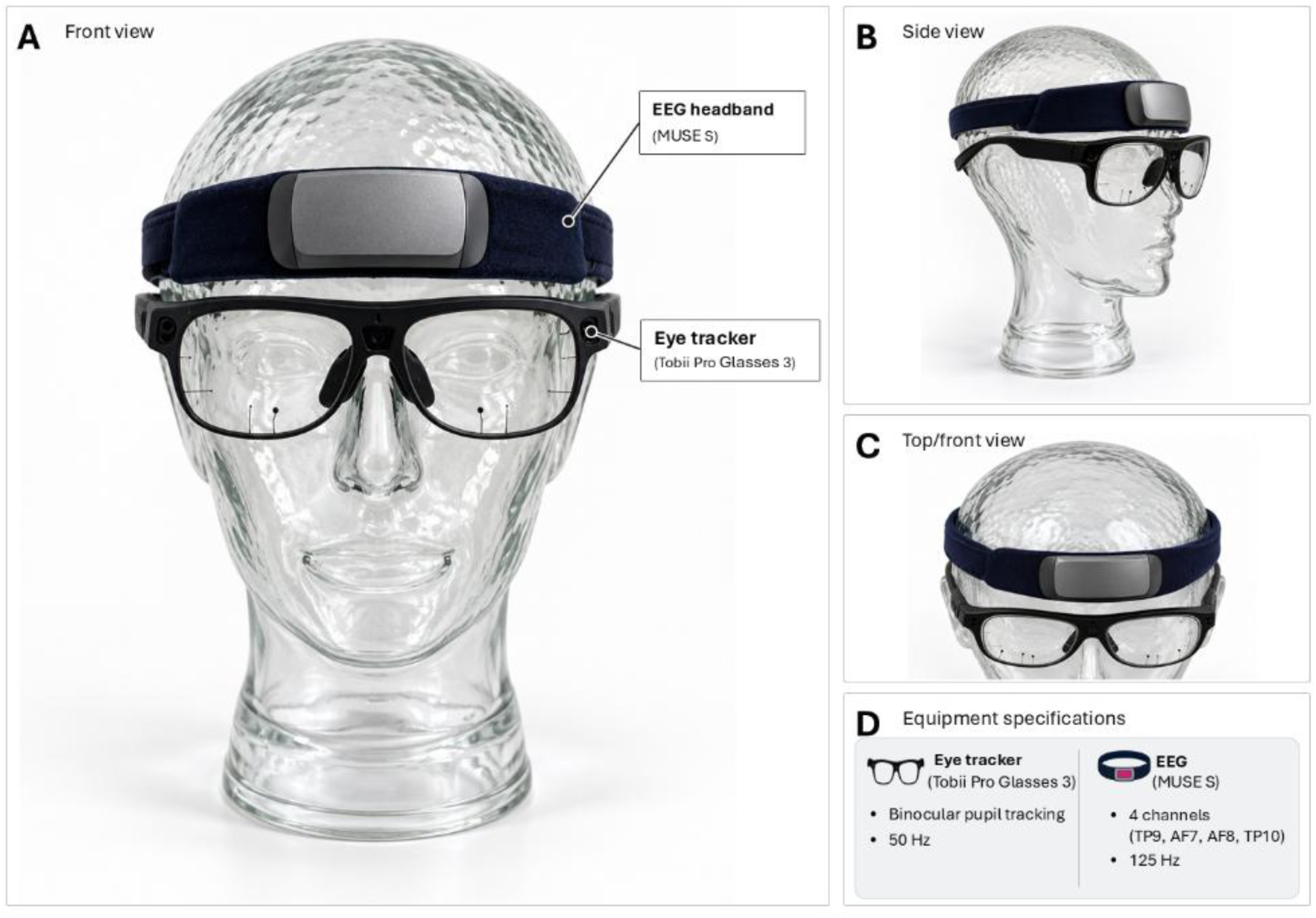
Wearable setup for pupil diameter and EEG recording. *Note.* (A-C) Frontal, side, and elevated frontal views of the Tobii Pro Glasses 3 eye tracker worn together with the MUSE S headband. (D) Acquisition specifications for binocular pupil diameter and four-channel EEG recordings.

Before each recording session, the system clocks of the recording devices were synchronized to a common network time reference to provide a shared absolute time reference across acquisition systems. Data from the MUSE and Polar devices were acquired concurrently through a custom acquisition application (developed by VirgoBit) that integrated the respective device streams and assigned a common timestamp to incoming data packets, as described in Criscuolo et al. (2026). Pupil data were recorded separately using the Tobii Pro Glasses 3 Controller App (Tobii AB, Danderyd, Sweden; https://www.tobii.com/products/eye-trackers/wearables/tobii-pro-glasses-3/controller-app).

### 2.4 Data preprocessing

Temporal synchronization across independent wearable recording systems is challenging because different data streams may have distinct sampling rates and timing references. As the EEG/heart rate and eye-tracking systems operated as separate recording streams, the recordings were temporally synchronized using their absolute timestamps. Only the overlapping time interval shared across the simultaneously recorded signals was retained for subsequent processing and analysis. The signals were then aligned to a common time axis to enable sample-wise correspondence between EEG and pupil recordings. Further details on multimodal data integration and synchronization framework are provided in Criscuolo et al. (2026).

#### 2.4.1 Pupil preprocessing

Pupil-data quality was first assessed separately for each condition based on the proportion of missing samples for each eye and the correlation between the left- and right-eye pupil traces. Inter-eye correlation was used as an additional quality-control measure, given that pupil fluctuations are generally correlated between eyes (Klingner et al., 2008; Kret & Sjak-Shie, 2019). Participants were excluded when more than 30% of data from both eyes were missing. When data from only one eye exceeded this threshold, data from the other eye was retained. When data from both eyes met the missing-data criterion but the correlation was below .7, the one with better data quality was retained. Otherwise, both were included in the subsequent preprocessing. Based on these criteria, two participants were excluded because data from both eyes contained at least 30% missing values. Single-eye data were used for three participants during resting state and for four participants during interoceptive attention.

Blink-related artifacts were identified using complementary missing data- and velocity-based criteria, consistent with established approaches for pupil-data preprocessing (Hershman et al., 2018; Kret & Sjak-Shie, 2019). Missing data segments lasting at least 60 ms were considered blink candidates. For binocular recordings, missing segments for the two eyes were required to overlap or occur within 50 ms of one another. Rapid pupil-size changes that were not represented as missing data were additionally identified using a robust velocity-based criterion. Pupil velocity was calculated as the absolute first-order change in pupil diameter, and artifacts were identified relative to the median and median absolute deviation of the velocity distribution (Kret & Sjak-Shie, 2019). For binocular recordings, velocity-based artifacts were retained only when corresponding events occurred in both eyes within the same 50 ms tolerance window. Missing data- and velocity-based candidates were combined, and events separated by less than 250 ms were merged. The resulting artifact windows were extended by 50 ms before onset and 200 ms after offset. Across participants, these blink- and artifact-related windows accounted on average for 13.92% of the recording during resting state and for 13.48% during interoceptive attention. Blink- and artifact-related missing segments of up to 3 s were reconstructed separately for each eye using piecewise cubic Hermite interpolation (PCHIP), consistent with common approaches for reconstructing short missing intervals in pupillometry data (Kret & Sjak-Shie, 2019; Mathôt et al., 2018). Remaining artifacts were subsequently identified using robust derivative- and amplitude-based criteria and reconstructed using PCHIP interpolation. This additional artifact screening affected on average 4.90% and 4.42% of the recordings during resting state and interoceptive attention, respectively.

The cleaned pupil traces were band-pass filtered between .005 and 2 Hz using a Butterworth IIR filter with zero-phase forward–backward filtering. For binocular recordings, the cleaned and filtered left- and right-eye traces were subsequently averaged sample-wise to obtain a single pupil time series. For recordings in which data from only one eye met the quality criteria, data from the retained eye was used as the final pupil trace.

#### 2.4.2 EEG preprocessing

EEG data (channels TP9, AF7, AF8, and TP10) were first screened for missing samples and flat line segments. Flat line segments were identified using a 100 ms sliding window with a standard deviation threshold of 10^−10^. Missing data gaps were reconstructed according to their duration. Gaps of up to 50 ms were interpolated using piecewise cubic Hermite interpolation (PCHIP, Fritsch & Carlson, 1980), gaps of 51-150 ms using Kalman smoothing (Rauch et al., 1965), and gaps of 151-1000 ms using forward-backward autoregressive interpolation (Kazlauskas & Pupeikis, 2014). Gaps exceeding 1 s were retained as missing. The EEG signals were subsequently high-pass filtered at 0.5 Hz using zero-phase filtering. Motion-related artifacts were attenuated using recursive least-squares (RLS) adaptive filtering based on the recorded inertial signals and their first- and second-order temporal derivatives (Masterton et al., 2007; Sweeney et al., 2012). Cardiac events were identified using the simultaneously recorded ECG signal, while corresponding cardiac artifacts in the EEG were attenuated using an optimal basis set (OBS) approach (Niazy et al., 2005). Six participants were excluded because no usable ECG recording was available for reliable cardiac-event detection. A discrete Fourier transform (DFT) filter was used to attenuate line-related noise at 25, 50 and 100 Hz.

Remaining transient artifacts were first identified using a kurtosis-based procedure. Excess kurtosis was calculated in 200 ms sliding windows advanced in 10 ms steps, and windows exceeding a threshold of 5.0 were identified as artifactual. A 100 ms interval before and after each detected event was marked as missing and reconstructed using the duration-dependent interpolation procedure described above. Additional artifact screening was then performed sequentially using a fixed raw amplitude threshold of ±100 μV, a derivative-based criterion, and a normalized-amplitude criterion. Samples identified at each of these stages were extended by 25 ms on either side and reconstructed using modified Akima cubic Hermite interpolation (MAKIMA). Finally, residual sharp spikes were detected separately within each EEG channel using robust amplitude and first-derivative z scores based on the channel median and median absolute deviation. The resulting artifact-related gaps were reconstructed using the duration-dependent procedure described above, applied separately to each channel.

Following artifact detection and reconstruction, EEG data quality was assessed based on the proportion of samples remaining missing after preprocessing. Participants were excluded if more than 15% of samples remained missing in any EEG channel in either condition. Based on this criterion, two participants were excluded from subsequent analyses. Among the retained participants, transient artifacts affected on average 1.29% of samples during resting state and 0.84% during interoceptive attention. Additional amplitude- and derivative-based screening affected 3.23% and 2.79% of samples, respectively, while residual artifacts were identified in 2.29% and 2.20% of samples. After preprocessing, an average of .19% and .18% of samples remained missing during resting state and interoceptive attention, respectively.

#### 2.4.3 Time-frequency decomposition

Frequency-specific EEG power time courses were derived separately for each of the four EEG channels using complex Morlet wavelets as implemented in the FieldTrip toolbox (Oostenveld et al., 2011). Power was calculated at 30 logarithmically spaced frequencies between 2 and 125 Hz, consistent with the logarithmic organization of neural oscillatory frequencies (Buzsáki & Mizuseki, 2014), and the broad frequency-resolved approach used in previous pupil–brain coupling work (Pfeffer et al., 2022). To balance spectrotemporal resolution across this frequency range (Tallon-Baudry & Bertrand, 1999), three-cycle wavelets were used for frequencies up to 5 Hz, and seven-cycle wavelets were used for frequencies above 5 Hz. The resulting power time courses were subsequently used in cross-correlation and mutual information analyses.

### 2.5 Statistical analyses

Our first goal was to assess the feasibility of examining pupil–brain coupling via wearable devices. Thus, the cross-correlation and mutual information analysis steps detailed below followed the procedure employed by Pfeffer et al. (2022). The original analysis pipeline was adapted to specifics of the instrumentation employed. To further characterize directional coupling, we expanded to include directionality analyses via Granger causality.

#### 2.5.1 Cross-correlation analysis

To characterize the temporal relationship between spontaneous pupil fluctuations and frequency-specific EEG activity, the power time courses derived from the four EEG channels were averaged at each frequency. Cross-correlations were then computed between pupil diameter and the channel-averaged EEG power time course separately for each participant and frequency across lags ranging from −10 to 10 s. Similar cross-correlation approaches were previously used to characterize temporal relationships between spontaneous pupil fluctuations and electrophysiological activity (Montefusco-Siegmund et al., 2022; Podvalny et al., 2021), including frequency-specific lead-lag relationships (Pfeffer et al., 2022). Positive lags indicate that EEG power fluctuations preceded pupil fluctuations, whereas negative lags indicate that pupil fluctuations preceded EEG power. For descriptive visualization, cross-correlations were additionally averaged within five frequency ranges: 2-4, 4-8, 8-12, 12-30, and 30-125 Hz. These ranges broadly correspond to conventional delta, theta, alpha, and beta bands, with frequencies above 30 Hz summarized as a single higher-frequency range. The band-averaged results were used only for visualization and descriptive summary; all statistical inferences were performed at the individual frequency level.

To determine whether the observed cross-correlations were higher than by chance, we used shuffling to generate a null distribution. For each participant, the pupil time series was divided into non-overlapping 20 s blocks. The blocks were then randomly reordered to disrupt the temporal alignment between pupil fluctuations and EEG power. Cross-correlations were recomputed for each shuffled time series, and the procedure was repeated 200 times.

Following the procedure employed by Pfeffer et al. (2022), at each frequency and lag, the observed cross-correlation coefficient was standardized relative to the corresponding null distribution by first subtracting the null mean and then dividing by the null standard deviation. The resulting Z-score maps were entered into group-level statistical analyses.

To identify frequencies at which the observed cross-correlations significantly exceeded the corresponding null distributions, a one-sided cluster-based permutation test against zero was conducted separately at each frequency within a 2 s lag window (Maris & Oostenveld, 2007). Adjacent lag points exceeding a cluster-forming threshold of *p* < .05 were grouped into clusters, and cluster-level significance was evaluated using 10,000 permutations. For frequencies containing a significant cluster at positive lags, the lag corresponding to the maximum group-mean Z-score within the cluster was identified. The representative lag was defined as the mean of these peak-lag estimates across the selected frequencies.

#### 2.5.2 Mutual information analysis

The cross-correlation analysis primarily characterizes the temporal structure of the association between EEG power and pupil fluctuations. We additionally used mutual information to quantify the strength of statistical dependence between frequency-specific EEG power and pupil diameter. Unlike cross-correlation, mutual information is not restricted to linear dependencies and can capture more general relationships between the two signals (Ince et al., 2017). As in Pfeffer et al. (2022), before mutual information estimation, the signals were temporally aligned based on the representative positive lags identified in the cross-correlation analysis. The representative lags were 752 ms during resting state and 777 ms during interoceptive attention. To apply a common temporal alignment across conditions, we used the mean lag of 764 ms, pairing EEG power with subsequent pupil diameter accordingly.

Mutual information was estimated using the Gaussian-copula mutual information method implemented in the GCMI toolbox (Ince et al., 2017). For each condition, participant, and frequency, the power time courses from the four EEG channels were jointly entered as a multivariate EEG variable. Mutual information was estimated between this multivariate EEG representation and the lag-aligned pupil signal across the entire recording. This procedure yielded one frequency-resolved mutual information spectrum for each participant and condition.

Participant-specific null distributions were generated using the same block-shuffling procedure described for the cross-correlation analysis. Mutual information was recalculated for 200 shuffled pupil time series using the same analysis pipeline applied to the original, unshuffled data. Observed mutual information estimates and their corresponding null distributions were log-transformed before the observed values were standardized relative to the null distributions. Positive Z scores indicated stronger pupil-EEG dependence than expected under the block-shuffled null model. To identify contiguous frequency ranges showing above-null mutual information, Z scores were tested against zero using a one-sided cluster-based permutation test across adjacent frequencies. Adjacent frequencies exceeding the cluster-forming threshold of *p* < .05 were grouped into clusters, and cluster-level significance was evaluated using 10,000 permutations.

#### 2.5.3 Spectral Granger causality analysis

Directional dependencies between EEG and pupil dynamics were examined using spectral Granger causality. Granger causality assesses whether the past values of one time series improve the prediction of another time series beyond the information contained in its own past (Granger, 1969). Its spectral formulation allows such directional coupling to be characterized as a function of frequency (Geweke, 1982).

The analysis was performed on the preprocessed and temporally synchronized EEG and pupil time-domain signals. The four EEG channels and the pupil signal were linearly detrended and then standardized separately within each channel using the mean and standard deviation of the finite samples. Following standardization, the four EEG channels were averaged sample-wise to obtain a single mean EEG time series. The continuous recordings were divided into 30 s overlapping windows with a step size of 10 s. A multivariate autoregressive (MVAR) model was fitted jointly to the mean EEG and pupil time series using the BSMART method implemented in FieldTrip (Cui et al., 2008; Oostenveld et al., 2011). Model order was evaluated separately for each participant and condition across candidate orders up to 250 samples using the Akaike information criterion (AIC) and Bayesian information criterion (BIC). Based on these model order estimates, we selected a fixed model order of 200 samples, which was consistently applied across participants and conditions to ensure a uniform model specification. At 250 Hz, this corresponded to a temporal history of 0.8 s. The resulting Granger-causality spectra were averaged across windows within each participant to obtain a participant-level spectrum for each direction.

The fitted MVAR model was converted into a spectral representation, from which frequency-domain Granger causality was computed using FieldTrip (Oostenveld et al., 2011), following Geweke’s formulation (Geweke, 1982). Estimates were obtained at 1 Hz intervals between 1 and 80 Hz in both the EEG-to-pupil and pupil-to-EEG directions. Given the sampling rate of 250 Hz, the Nyquist frequency was 125 Hz. The upper frequency limit was set below the Nyquist frequency to reduce potential instability in autoregressive spectral estimates near this boundary (Richter et al., 2018). Null distributions were generated using the same block-shuffling procedure described for the cross-correlation analysis. For each shuffled time series, Granger causality was recalculated across the same 30 s windows, and the resulting spectra were averaged across windows. This procedure was repeated 200 times, yielding a null distribution of participant-level Granger-causality spectra for each direction. At each frequency and for each direction, the observed Granger causality estimate was standardized relative to the corresponding surrogate distribution by subtracting the surrogate mean and dividing the surrogate standard deviation, yielding a Z score.

## 3. Results

### 3.1 Resting-state pupil–brain coupling

We first characterized resting-state pupil–brain coupling in terms of its temporal, frequency-dependent, and directional organization using cross-correlation, mutual information, and spectral Granger causality, respectively.

#### 3.1.1 Temporal organization of resting-state pupil–brain coupling

Cross-correlation profiles differed across frequency ranges (Fig. 2 A, B). Within the 2-12 Hz range, correlation maxima occurred at positive lags, indicating that EEG power fluctuations preceded subsequent pupil fluctuations. The maximum correlations occurred at approximately 472, 600, and 628 ms for the 2-4, 4-8, and 8-12 Hz bands, respectively. In contrast, the 12-30 and 30-125 Hz bands did not exhibit clear correlation peaks or a consistent lag structure (Fig. 2B).

**Fig. 2.**
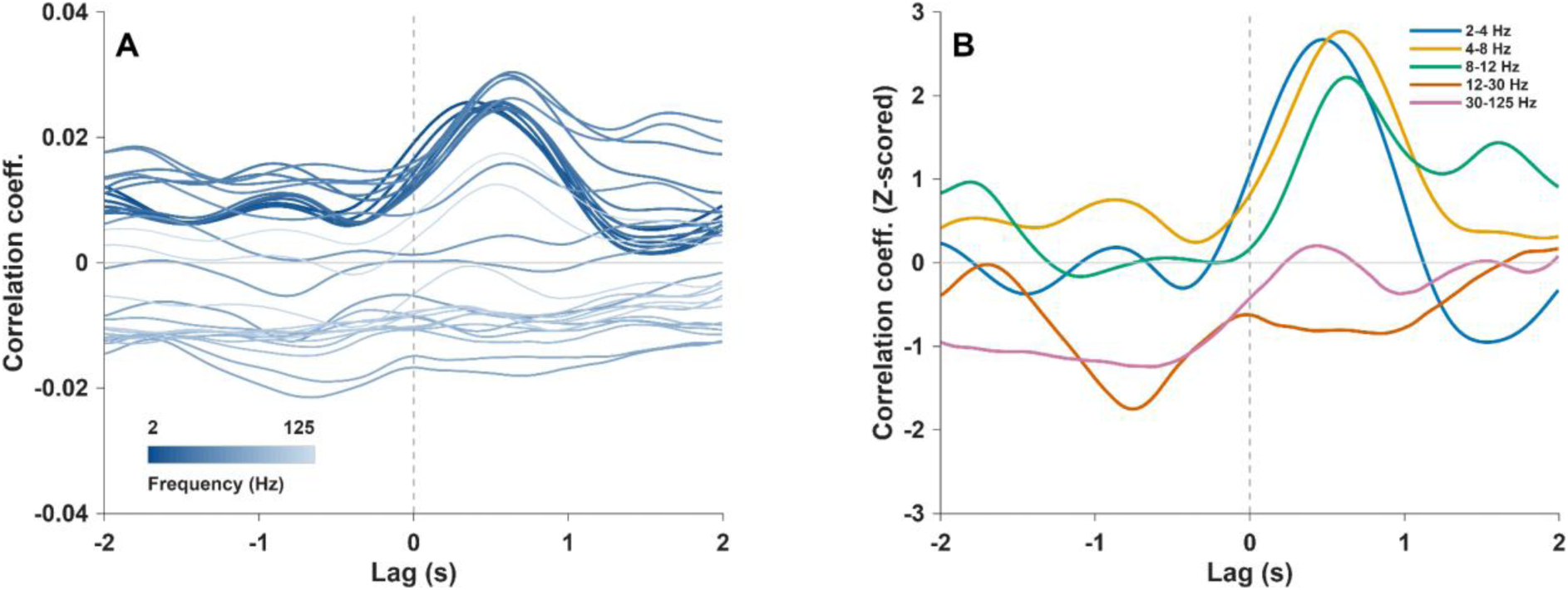
Frequency-resolved cross-correlation between EEG power and pupil diameter during resting state. *Note.* (A) Lines show group-mean cross-correlation coefficients (y-axis) across 30 logarithmically spaced frequencies from 2 to 125 Hz, and with temporal lags ranging from −2 to +2 s (x-axis). Darker lines represent lower frequencies, and lighter lines represent higher frequencies. (B) Lines show group-mean cross-correlation profiles averaged within the 2-4, 4-8, 8-12, 12-30, and 30-125 Hz frequency bands. Positive lags indicate that EEG power fluctuations preceded pupil fluctuations, whereas negative lags indicate that pupil fluctuations preceded EEG power fluctuations. The vertical dashed line marks zero lag.

To obtain a representative estimate of the temporal offset, the observed cross-correlations were further standardized relative to participant-specific null distributions. Frequency-wise cluster permutation tests identified positive-lag clusters reaching the nominal cluster-level threshold of *p* < .05 at several frequencies. Averaging the peak-lag estimates yielded a representative resting-state lag of 752 ms.

#### 3.1.2 Frequency-dependent statistical dependence

Following temporal alignment using a lag of approximately 764 ms, calculated by averaging the two condition mean lags from the cross-correlation analyses (752 ms in resting state and 777 ms in interoceptive attention), we examined pupil–brain coupling via mutual information.

Pupil-EEG mutual information exceeded the null distribution across two frequency ranges (Fig. 3A). Standardized mutual information was significantly greater than zero across 2.66-11.07 Hz (*p*cluster = .004) and 34.64-81.49 Hz (*p*cluster = .022). Dots in Fig. 3A indicate frequencies belonging to clusters that survived cluster-level correction. Thus, resting-state pupil-EEG dependence was evident across lower frequencies, with an additional higher-frequency component.

**Fig. 3.**
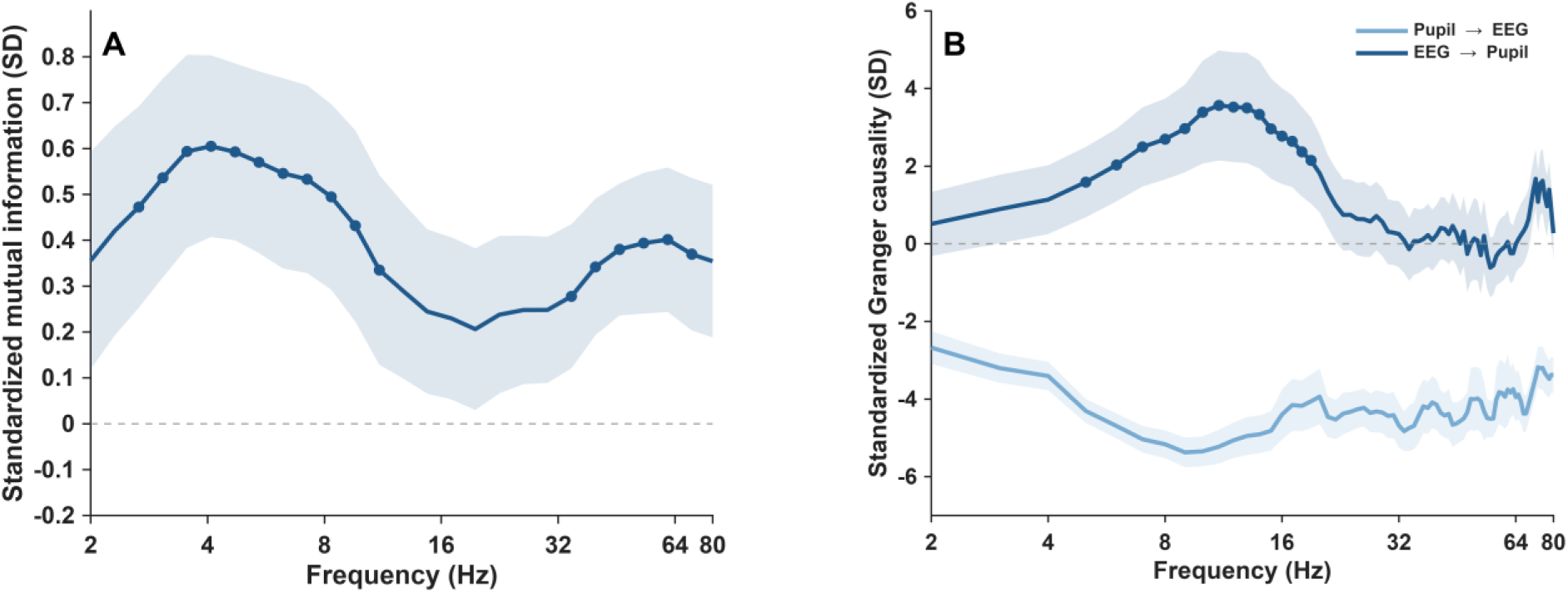
(A) Frequency-resolved mutual information between EEG power and pupil diameter during resting state. (B) Frequency-resolved spectral Granger causality between EEG activity and pupil diameter during resting state. *Note.* (A) The line shows group-mean mutual-information Z scores, and the shaded area indicates ±1 SD across participants. Estimates were standardized relative to null distributions generated from 200 block-shuffled pupil time series; positive Z scores indicate stronger pupil-EEG dependence than expected under the null model. Filled circles denote frequencies belonging to significant positive clusters. Frequencies are displayed on a logarithmic scale from 2 to 80 Hz. (B) Lines show group-mean spectral Granger-causality Z scores for the pupil-to-EEG and EEG-to-pupil directions, and shaded areas indicate ±1 SD across participants. Estimates were standardized relative to null distributions generated from 200 block-shuffled pupil time series; positive Z scores indicate stronger directional coupling than expected under the null model. Filled circles denote frequencies belonging to significant positive clusters identified by cluster-based permutation tests. Frequencies are displayed on a logarithmic scale from 2 to 80 Hz.

#### 3.1.3 Directional organization of pupil–brain coupling in resting-state

During resting state, EEG-to-pupil Granger causality significantly exceeded the null distribution across 5-19 Hz (*p*cluster = .011), whereas pupil-to-EEG Granger causality showed no significant above-null clusters (Fig. 3B). Direct comparison of the two directions revealed significantly stronger EEG-to-pupil than pupil-to-EEG Granger causality across 3-49 Hz (*p*cluster < .001), 51-58 Hz (*p*cluster = .010), 60-74 Hz (*p*cluster = .002), and 76-80 Hz (*p*cluster = .016). These results indicate that directional coupling in resting state was predominantly organized from EEG to pupil, with above-null effects mainly concentrated at lower frequencies.

### 3.2 Pupil–brain coupling during interoceptive attention

We next examined whether the overall spectrotemporal organization of pupil–brain coupling observed at rest was also present during interoceptive attention. As during resting state, cross-correlation maxima within the 2-12 Hz range occurred at positive lags (Supplementary Fig. S1 A, B), indicating that EEG power fluctuations preceded subsequent pupil fluctuations. Peak correlations occurred at approximately 416, 572, and 708 ms for the 2-4, 4-8, and 8-12 Hz bands, respectively. At higher frequencies, the 12-30 and 30-125 Hz bands showed later and broader positive-lag maxima at approximately 1180 and 1924 ms, respectively. Frequency-wise cluster permutation tests identified positive-lag clusters reaching the nominal cluster-level threshold of *p* < .05 at several frequencies, with a mean peak lag of 777 ms across these frequencies.

Mutual information also exceeded the null distribution at lower frequencies during interoceptive attention (Supplementary Fig. S2). Significant clusters were identified across 2.00-3.54 Hz (*p*cluster = .035) and 4.71-8.32 Hz (*p*cluster = .042). In contrast to the resting-state analyses, there was no additional significant higher-frequency cluster.

Directional coupling was also evident during interoceptive attention. EEG-to-pupil Granger causality exceeded the null distribution across 8-13 Hz (*p*cluster = .033), whereas pupil-to-EEG Granger causality showed no significant above-null clusters (Supplementary Fig. S2B). Direct comparison of the two directions revealed significantly stronger EEG-to-pupil than pupil-to-EEG Granger causality across 2-29 Hz, 31-49 Hz, 51-74 Hz (all *p*cluster < .001), and 76-80 Hz (*p*cluster = .012).

## 4. Discussion

The present study examined whether the spectrotemporal and directional organization of spontaneous pupil–brain coupling can be captured using lightweight and principally mobile EEG and eye tracking. Resting-state pupil–brain coupling showed a clear frequency-dependent organization, with lower EEG frequencies showing the clearest convergence across analyses, alongside additional effects at higher frequencies. EEG power fluctuations preceded subsequent pupil fluctuations by approximately 760 ms, and directional coupling was predominantly organized from EEG to pupil across low-to-mid frequencies. In the resting state, but not during interoceptive attention, we also observed additional higher-frequency coupling. Besides this difference, a broadly similar core organization was observed during interoceptive attention. Together, these findings confirm that the mobile recording setup successfully captured key features of spontaneous pupil–brain coupling previously observed using stationary systems.

Our findings are broadly consistent with previous studies, particularly regarding the spectrotemporal organization of pupil–brain coupling. Specifically, EEG power fluctuations generally preceded changes in pupil diameter, with the clearest temporal relationships observed at low frequencies (Montefusco-Siegmund et al., 2022; Pfeffer et al., 2022). At the spectral level, we observed significant statistical dependence between pupil diameter and EEG power across the delta-to-alpha frequency range. The observed delay is also compatible with the physiological dynamics of pupil control. Pupil diameter is a peripheral autonomic output, regulated through sympathetic and parasympathetic pathways acting on the iris musculature (McDougal & Gamlin, 2015), and pupillary responses evolve over several hundred milliseconds following changes in their underlying input (Hoeks & Levelt, 1993). The convergence of these two analyses indicates that lower-frequency EEG dynamics are particularly involved in spontaneous pupil–brain coupling and may play a role in the neuromodulatory processes influencing pupil diameter (Reimer et al., 2016). This pattern is like previous work linking pupil-related arousal to changes in cortical state. In awake rodents, spontaneous pupil fluctuations closely track transitions between synchronized and desynchronized cortical states, which are prominently reflected in approximately 2-10 Hz cortical activity (McGinley et al., 2015; Reimer et al., 2016). Pupil dilation was associated with reduced low-frequency activity and cortical desynchronization, whereas constriction was accompanied by stronger low-frequency fluctuations (Reimer et al., 2014). Lower-frequency EEG activity may therefore be particularly sensitive to spontaneous fluctuations in cortical states that covary with pupil-linked arousal.

However, the specific spectral relationships observed during resting state were not fully consistent with previous human studies. Pfeffer et al. (2022) reported a negative relationship between pupil diameter and cortical power in the delta range, whereas correlations over a similar low-frequency range were predominantly positive in the present study. The spectral distribution of mutual information also differed. Pfeffer et al. (2022) reported the strongest MI effects at intermediate frequencies, together with additional dependence at lower and higher frequencies. In the present study, dependence was most evident at lower frequencies, with an additional significant cluster in the gamma range. These differences may partly reflect distinct characteristics in spatial coverage and analytical approaches. Their source-level analyses showed that pupil–brain coupling was not spatially uniform. Low-frequency relationships were relatively widespread, but intermediate- and high-frequency coupling showed more region-specific patterns, including associations of different directions across cortical areas (Pfeffer et al., 2022). Several human EEG and MEG studies have nevertheless reported partly similar pupil–brain relationships. Ceh et al. (2020) and Montefusco-Siegmund et al. (2022) found positive pupil-related associations with alpha activity, particularly over posterior scalp regions, while Podvalny et al. (2021) reported pupil-linked cortical effects across both lower and higher frequencies across large-scale cortical networks. The EEG system used in the present study covered only frontal and temporal scalp sites, potentially capturing a different combination of cortical activity. Furthermore, we estimated mutual information after temporally aligning EEG power and pupil fluctuations based on the cross-correlation results. David et al. (2025) showed that the direction of low-frequency pupil-EEG associations can vary as a function of temporal lag. Thus, these findings support a broad correspondence with previous studies rather than an exact replication of their frequency-specific coupling profiles.

Beyond its temporal and spectral relationships identified from frequency-specific EEG power, spectral Granger-causality analysis provided a complementary characterization of directional coupling between the EEG and pupil signals. EEG-to-pupil Granger causality significantly exceeded the null distribution at lower frequencies, whereas pupil-to-EEG Granger causality showed no significant effects. This finding complements the low-frequency coupling identified by cross-correlation and mutual information, while providing additional information about the directional organization. Direct comparison of the two directions further showed a broader predominance of EEG-to-pupil over pupil-to-EEG. Therefore, these results suggest that the directional organization of spontaneous pupil–brain coupling was predominantly characterized by EEG-to-pupil predictability, with the clearest phenomenon at lower frequencies.

One possible physiological interpretation is that spontaneous cortical dynamics contribute to subsequent changes in pupil diameter through interactions with neuromodulatory and autonomic systems. Cortical regions involved in arousal and autonomic regulation, including prefrontal, cingulate, and insular areas, can influence locus coeruleus circuitry, while frontal regions are connected to the superior colliculus. These pathways provide plausible routes through which cortical activity may modulate pupillary control (Arnsten & Goldman-Rakic, 1984; Joshi et al., 2016; Peinkhofer et al., 2019). However, interactions between cortical and neuromodulatory systems are not unidirectional. Phasic locus coeruleus activity precedes shifts in prefrontal gamma activity (Neves et al., 2018). Conversely, transient prefrontal high-gamma signals predict later changes in locus coeruleus population activity (Totah et al., 2021). Alternatively, cortical activity may reflect changes in neuromodulatory and autonomic systems earlier than these changes become apparent in pupil diameter. Concurrent recordings have shown that locus coeruleus spiking precedes both cortical activity and pupil dilation, but with different temporal delays (Yang et al., 2021). In this case, EEG activity may temporally precede pupil fluctuations without directly contributing to their generation. Moreover, Granger causality establishes directional coupling rather than causal influence (Barnett & Seth, 2014; Seth et al., 2015). The observed EEG-to-pupil predictability may reflect interactions within a broader cortical-neuromodulatory system, potentially including shared neural influences on cortical and pupil dynamics.

The secondary interoceptive attention condition provided an opportunity to examine whether the core organization identified during resting state remained detectable when attention was directed to bodily signals. Overall, resting state and interoceptive attention conditions showed consistent patterns. EEG activity again preceded subsequent pupil fluctuations at low frequencies, mutual information showed significant low-frequency pupil-EEG dependence, and directional coupling was predominantly organized from EEG to pupil across low frequencies. The main difference was observed at higher frequencies. Mutual information did not show a significant higher-frequency component. Cross-correlation also showed later and broader positive-lag maxima at higher frequencies, but these were descriptive features of the band-averaged profiles rather than robust inferential effects. Previous studies have shown that interoceptive attention can modulate frequency-specific cortical activity, including alpha and beta activity (Crivelli & Balconi, 2025; Pultsina et al., 2026), providing a possible context for the descriptive spectral differences observed here. However, the interoceptive attention condition simply required participants to focus on internal bodily signals, and the present study was not designed to directly test state-related differences. These findings therefore primarily support the preservation of the core spectrotemporal, and directional organization of pupil–brain coupling across states, rather than a specific effect of interoceptive attention.

## 5. Limitations

The present study has several limitations. First, the four-channel EEG system provided substantially more limited spatial sampling than conventional EEG or whole-head MEG. Compared to conventional gel EEG caps, the MUSE headband relies on dry electrodes, which can raise electrode-skin impedance and increase variability in signal quality. Preprocessing and quality control procedures for wearable EEG systems with a small number of channels such as MUSE are also less standardized and less extensively validated than those commonly used for conventional EEG recordings. The limited number of electrodes further restricts the use of procedures such as ICA-based artifact correction and spatial interpolation. These limitations constrain both the spatial interpretation of the present findings and the extent to which standard EEG preprocessing approaches can be directly applied. Second, the higher-frequency findings require further validation. Scalp EEG at these frequencies is particularly susceptible to contamination from cranial muscle and ocular activity (Hipp & Siegel, 2013; Muthukumaraswamy, 2013). Finally, although the EEG and eye tracking systems were lightweight and mobile, recordings were obtained under controlled laboratory conditions. Participants remained seated and followed relatively constrained task instructions, including visual fixation during resting state. It therefore remains to be tested whether the same spectrotemporal, and directional organization can be reliably detected during movement, natural visual exploration, or other everyday behaviors.

## 6. Implications and future directions

The present findings support the feasibility of characterizing spontaneous pupil–brain coupling using lightweight EEG and eye tracking. Beyond methodological feasibility, studying pupil–brain coupling may help clarify how spontaneous arousal fluctuations are coordinated with ongoing cortical activity (Liu et al., 2026). As pupil dynamics reflect the combined influence of multiple neuromodulatory and autonomic processes, their coupling with EEG may provide additional information about ongoing brain state fluctuations beyond pupil fluctuations. A future research direction is therefore to examine whether the spectrotemporal, and directional structure of pupil–brain coupling changes with fluctuations in different arousal, attentional engagement, and behavioral state. A more flexible recording setup could facilitate such experiments to less constrained behavioral contexts, including natural listening, visual exploration, and everyday behaviors. Future work should also test whether changes in pupil– brain coupling are associated with behavioral performance and whether these relationships remain stable across individuals and contexts. Together, these developments could help clarify whether pupil–brain coupling reflects ongoing brain-state transitions and establishes its functional relevance in more naturalistic environments.

## 7. Conclusions

In conclusion, spontaneous pupil–brain coupling during resting state showed a structured spectrotemporal and directional organization. EEG activity generally preceded subsequent pupil fluctuations, lower frequencies showed the clearest convergence across analyses, and directional coupling was predominantly organized from EEG to pupil across low frequencies. A broadly similar core organization was also observed during interoceptive attention. Importantly, these patterns could be captured using lightweight EEG and eye tracking, supporting the feasibility of investigating ongoing pupil–brain dynamics in more flexible and naturalistic recording settings.

## CRediT authorship contribution statement

**Tongxin Liu**: Writing – Review & Editing, Writing – Original draft, Methodology, Formal analysis, Data curation, Conceptualization. **Sonja A. Kotz**: Writing – Review & Editing, Supervision, Project administration, Methodology, Conceptualization. **Michael Schwartze**: Writing – Review & Editing, Supervision, Project administration, Methodology, Conceptualization. **Antonio Criscuolo**: Writing – Review & Editing, Supervision, Project administration, Methodology, Software, Data curation, Conceptualization.

## Acknowledgements

We thank all participants who participated in the study for their contribution, time, and effort.

## Data availability

Data will be made available on request.

