## Supplemental Data 1 for "Capturing continuous spectrotemporal pupil–brain coupling with simultaneous wearable EEG/Eye tracking"

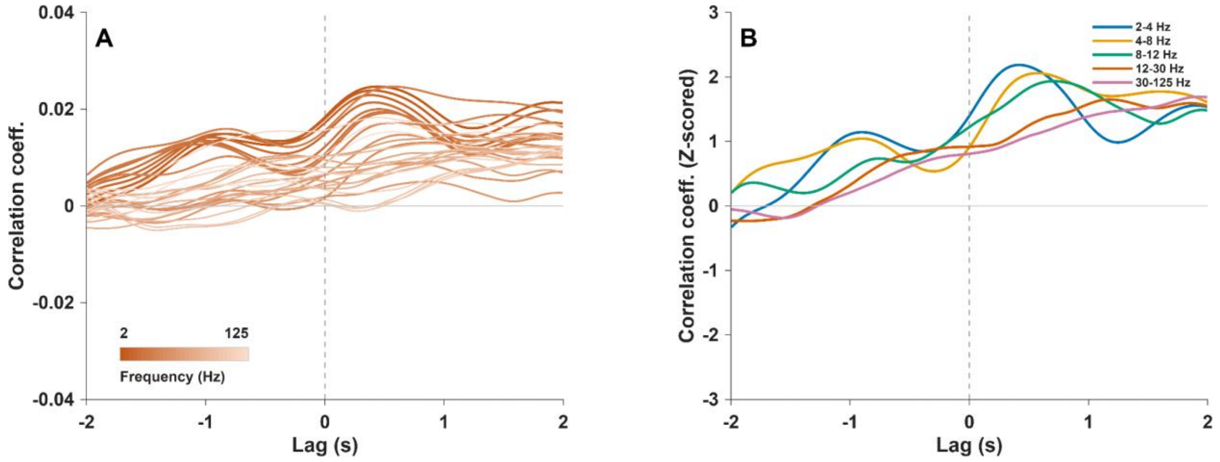


**Supplementary Fig. S1**. *Frequency-resolved cross-correlation between EEG power and pupil diameter during interoceptive attention.*

*Note.* (A) Lines show group-mean cross-correlation coefficients across 30 logarithmically spaced frequencies from 2 to 125 Hz. Darker lines represent lower frequencies, and lighter lines represent higher frequencies. (B) Lines show group-mean cross-correlation profiles averaged within the 2-4, 4-8, 8-12, 12-30, and 30-125 Hz frequency bands. For visualization, lags from −2 to 2 s are displayed. Positive lags indicate that EEG power fluctuations preceded pupil fluctuations, whereas negative lags indicate that pupil fluctuations preceded EEG power fluctuations. The vertical dashed line marks zero lag.

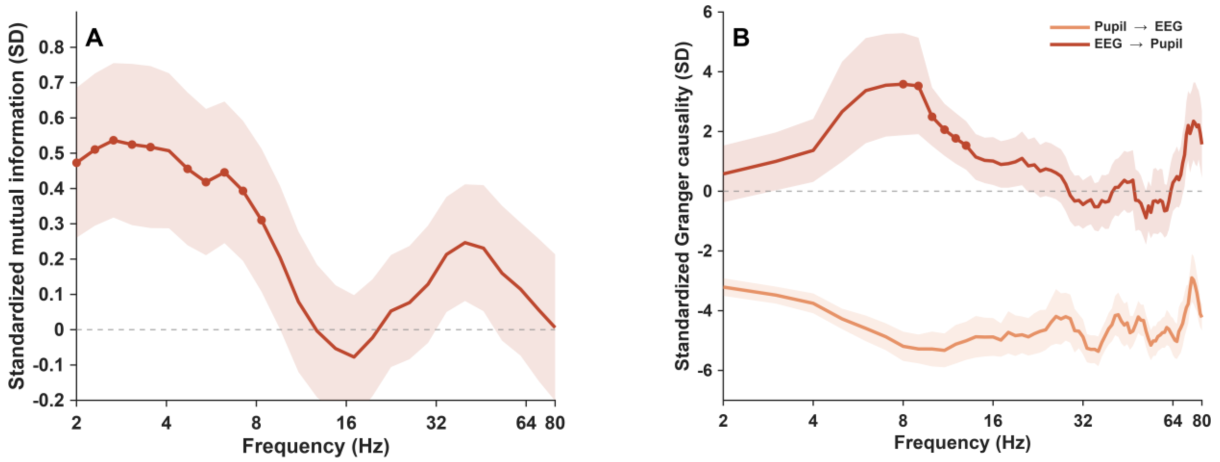


**Supplementary Fig. S2**. *(A) Frequency-resolved mutual information between EEG power and pupil diameter during interoceptive attention. (B) Frequency-resolved spectral Granger causality between EEG activity and pupil diameter during interoceptive attention.*

*Note.* (A) The line shows group-mean mutual-information Z scores, and the shaded area indicates ±1 SD across participants. Estimates were standardized relative to null distributions generated from 200 block-shuffled pupil time series; positive Z scores indicate stronger pupil-EEG dependence than expected under the null model. Filled circles denote frequencies belonging to significant positive clusters. Frequencies are displayed on a logarithmic scale from 2 to 80 Hz. (B) Lines show group-mean spectral Granger-causality Z scores for the pupil-to-EEG and EEG-to-pupil directions, and shaded areas indicate ±1 SD across participants. Estimates were standardized relative to null distributions generated from 200 block-shuffled pupil time series; positive Z scores indicate stronger directional coupling than expected under the null model. Filled circles denote frequencies belonging to significant positive clusters identified by cluster-based permutation tests. Frequencies are displayed on a logarithmic scale from 2 to 80 Hz.
